# Stimulation of cardiac fibroblast Piezo1 channels opposes myofibroblast differentiation and induces IL-6 secretion via Ca^2+^-mediated p38 MAP kinase activation

**DOI:** 10.1101/603456

**Authors:** Nicola M. Blythe, Vasili Stylianidis, Melanie J. Ludlow, Hamish T. J. Gilbert, Elizabeth L. Evans, Kevin Cuthbertson, Richard Foster, Joe Swift, Jing Li, Mark J. Drinkhill, Frans A. van Nieuwenhoven, Karen E. Porter, David J. Beech, Neil A. Turner

## Abstract

Piezo1 is a mechanosensitive cation channel with widespread physiological importance; however its role in the heart is poorly understood. Cardiac fibroblasts are responsible for preserving the structural integrity of the myocardium and play a key role in regulating its repair and remodeling following stress or injury. We investigated expression and function of Piezo1 in cultured human and mouse cardiac fibroblasts. RT-PCR studies confirmed expression of *Piezo1* mRNA in cardiac fibroblasts at similar levels to endothelial cells. Fura-2 intracellular Ca^2+^ measurements validated Piezo1 as a functional ion channel that was activated by the Piezo1 agonist, Yoda1. Yoda1-induced Ca^2+^ entry was inhibited by Piezo1 blockers (gadolinium, ruthenium red) and the Ca^2+^ response was reduced proportionally by Piezo1 siRNA knockdown or in cells from *Piezo1*^+/−^ mice. Investigation of Yoda1 effects on selected remodeling genes indicated that Piezo1 activation opposed cardiac fibroblast differentiation; data confirmed by functional collagen gel contraction assays. Piezo1 activation using Yoda1 or mechanical stretch also increased the expression of interleukin-6 (IL-6), a mechanosensitive pro-hypertrophic and pro-fibrotic cytokine, in a Piezo1-dependent manner. Multiplex kinase activity profiling combined with kinase inhibitor studies and phospho-specific western blotting, established that Piezo1 activation stimulated IL-6 secretion via a pathway involving p38 MAP kinase, downstream of Ca^2+^ entry. In summary, this study reveals that cardiac fibroblasts express functional Piezo1 channels coupled to reduced myofibroblast activation and increased secretion of paracrine signaling molecules that can modulate cardiac remodeling.

## Introduction

Cardiac fibroblasts play important roles in the normal physiology of the heart and in its response to damage or stress. Mechanical stretching of cardiac fibroblasts, that occurs secondary to cardiac dilatation or changes in hemodynamic burden in the heart, induces differentiation of fibroblasts into myofibroblasts with increased expression of pro-fibrotic cytokines, extracellular matrix (ECM) proteins and ECM receptors [1–2]. This preserves cardiac integrity and performance but the response can become maladaptive; surplus deposition of cardiac ECM results in fibrosis, which increases the stiffness of the myocardium and reduces pumping capacity [3].

Much remains unknown regarding how mechanical forces are translated into transcriptional responses important for determination of the fibroblast phenotype and how this can lead to cardiac remodeling [4,5]. Stretch-activated ion channels have been identified as candidates for cardiac mechanotransducers [6]. Intracellular Ca^2+^ controls many functions in cardiac fibroblasts, including ECM synthesis and cell proliferation [7,8] and it is known that Ca^2+^ entry through ion channels is important for cardiac fibroblast responsiveness. For example, TRPV4 channel-mediated Ca^2+^ signaling has been shown to play a role in mechanosensation and the differentiation of fibroblasts to myofibroblasts [9].

In 2010, the Piezo1 and Piezo2 proteins were identified as the long-sought molecular carriers of an excitatory mechanically-activated current found in many cell types [10]. Piezo1, a non-selective cation channel, can be activated by a synthetic small molecule named Yoda1 [11]; a useful tool for activating the channel without the need for mechanical stimulation. *Piezo1* mRNA has been detected in the murine heart whereas *Piezo2* mRNA was barely detectable [10]. Piezo1 has recently been shown to be expressed in cardiomyocytes and upregulated in heart failure [12]. However, there are currently no reports on the role of Piezo1 in cardiac fibroblasts. We hypothesized that Piezo1 plays an important role in cardiac fibroblast function by regulating Ca^2+^ entry and downstream signaling.

Our data provide evidence that Piezo1 acts as a functional Ca^2+^-permeable ion channel in murine and human cardiac fibroblasts, and that its activation by Yoda1 opposes cardiac myofibroblast differentiation and is coupled to secretion of interleukin (IL)-6, a cytokine that is important in the response to cardiac injury and hypertrophic remodeling. We further revealed that Piezo1-induced Ca^2+^ entry was coupled to IL-6 expression via activation of p38 MAP kinase.

## Results

### Piezo1 expression and activity in cardiac fibroblasts

Messenger RNA encoding Piezo1 was detected in cultured cardiac fibroblasts from both mouse and human hearts (**Fig. 1A,B**). In a comparison with endothelial cells, which are known to express high levels of this channel [13], *Piezo1* mRNA expression levels in murine cardiac fibroblasts were similar to those observed in murine pulmonary endothelial cells, and those in human cardiac fibroblasts were similar to those in human saphenous vein endothelial cells and human umbilical vein endothelial cells (HUVECs). Using a magnetic antibody cell separation (MACS) technique [14], we confirmed that Piezo1 was expressed in the fibroblast-enriched (*Col1a1/Col1a2/Ddr2/Pdgfra*-positive [14]) fraction of freshly isolated cells from mouse heart at about half the level of the endothelial cell-enriched (*Pecam*-*1*-positive [14]) fraction (**Fig. 1C**). *Piezo1* mRNA levels were 20 times higher in isolated cardiac fibroblasts than cardiomyocytes (**Fig. 1C**).

**Figure 1.**
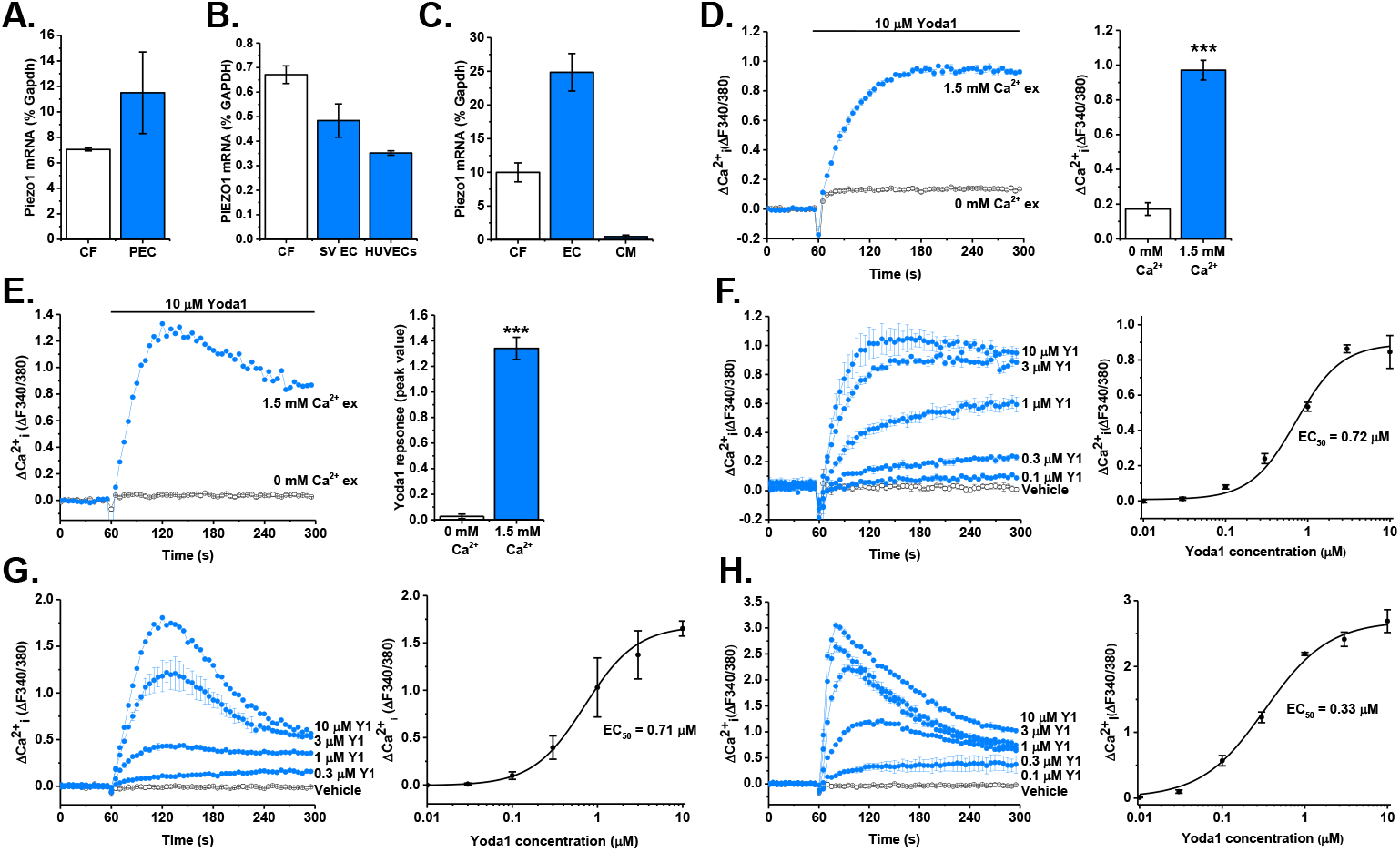
Piezo1 is expressed by cardiac fibroblasts and forms a functional ion channel. **(A,B)** RT-PCR analysis of *Piezo1* mRNA expression in (*A*) murine cardiac fibroblasts (CF; n=3) compared to murine pulmonary endothelial cells (PEC; n=2), and (*B*) human cardiac fibroblasts (CF; n=6) compared to human saphenous vein endothelial cells (SVEC, n=3) and human umbilical vein endothelial cells (HUVEC; n=3). Expression measured as % of housekeeping control (*Gapdh/GAPDH*). **(C)** RT-PCR analysis of *Piezo1* mRNA expression in the endothelial cell-enriched fraction 1 (EC) and the fibroblast-enriched fraction 2 (CF) isolated from murine heart using magnetic antibody cell separation (MACS) technique (n=4). Cardiomyocytes (CM) were isolated from separate hearts (n=3). Expression measured as % of housekeeping control (*Gapdh*). **(D,E)** Representative Ca^2+^ traces and mean ± SEM data are shown. Ca^2+^ entry evoked by 10 μM Yoda1 in murine (*D*) and human (*E*) cardiac fibroblasts in the presence or absence of extracellular Ca^2+^. ***P<0.001 (n/N=3/9). **(F-H)** Ca^2+^ entry evoked by varying concentrations of Yoda1 application at 60 sec, ranging from 0.1-10 μM in murine (*F*) and human (*G*) cardiac fibroblasts and HEK T-Rex-293 cells heterologously expressing mouse Piezo1 (*H*). Vehicle control is illustrated by the black trace. Mean ± SEM data are displayed as concentration-response curves and fitted curves are plotted using a Hill Equation indicating the 50 % maximum effect (EC_50_) of Yoda1 (n/N=3/9).

Having demonstrated that cardiac fibroblasts express *Piezo1* mRNA, we investigated whether the Piezo1 protein was able to form a functional ion channel. Using the Fura-2 Ca^2+^ indicator assay, it was found that Yoda1, a Piezo1 agonist [11], elicited an increase in intracellular Ca^2+^ in murine and human cardiac fibroblasts (**Fig. 1D,E**). Consistent with the Yoda1-induced increase in intracellular Ca^2+^ being due to influx of extracellular Ca^2+^ through an ion channel, the Ca^2+^ signal was reduced by >90% when extracellular Ca^2+^ was absent in human and mouse cardiac fibroblast cultures (**Fig. 1D,E**).

Concentration-response data for Yoda1 in murine cardiac fibroblasts revealed a marked effect at 0.3 μM and the maximal response was generated at 10 μM; the 50% maximum effect (EC_50_) of Yoda1 was estimated to be 0.72 μM (**Fig. 1F**). This was almost identical to the EC_50_ observed in human cardiac fibroblasts in similar experiments (**Fig. 1G**) and comparable with those for mouse Piezo1 heterologously expressed in HEK T-REx™-293 cells, where the EC_50_ of Yoda1 was 0.33 μM (**Fig. 1H**).

Gadolinium (Gd^3+^) and ruthenium red, both non-specific inhibitors of mechanosensitive ion channels including Piezo1 [10], were used to investigate the pharmacology of the channel. Murine cardiac fibroblasts pre-incubated with these inhibitors exhibited significantly reduced Yoda1-evoked Ca^2+^ entry (>70% reduction in both cases) (**Fig. 2A**). Additionally, the Yoda1-evoked Ca^2+^ entry was significantly reduced by >50% by Dooku1 (**Fig. 2A**), an analogue of Yoda1 that has antagonist properties against Yoda1 in Piezo1-overexpressing HEK-293 cells and HUVECs [15]. The data were similar in human cardiac fibroblasts, where again all three inhibitors significantly reduced the Ca^2+^ influx in response to Yoda1 (**Suppl. Fig. 1**). The Ca^2+^ entry elicited by Yoda1 application was also investigated in cardiac fibroblasts isolated from a global heterozygous (Het) *Piezo1*^+/−^ mouse line. RT-PCR analysis confirmed the predicted 50% reduction in *Piezo1* mRNA expression in cardiac fibroblasts derived from *Piezo1*^+/−^ hearts compared with WT (**Fig. 2B**). The Ca^2+^ response to Yoda1 was reduced by 40% (**Fig. 2C**), whereas the response to ATP was similar in cells from WT and *Piezo1*^+/−^ mice (**Fig. 2D**). Piezo1-specific siRNA, which decreased *Piezo1* mRNA expression by 80% in murine cardiac fibroblasts (**Fig. 2E**), reduced Yoda1-evoked Ca^2+^ entry by a similar level (**Fig. 2F**), whereas control siRNA was without effect. Similar results were obtained with human cardiac fibroblasts (**Fig. 2G,H**). Thus, the Yoda1 response was dependent upon Piezo1 and proportional to its expression level.

**Figure 2.**
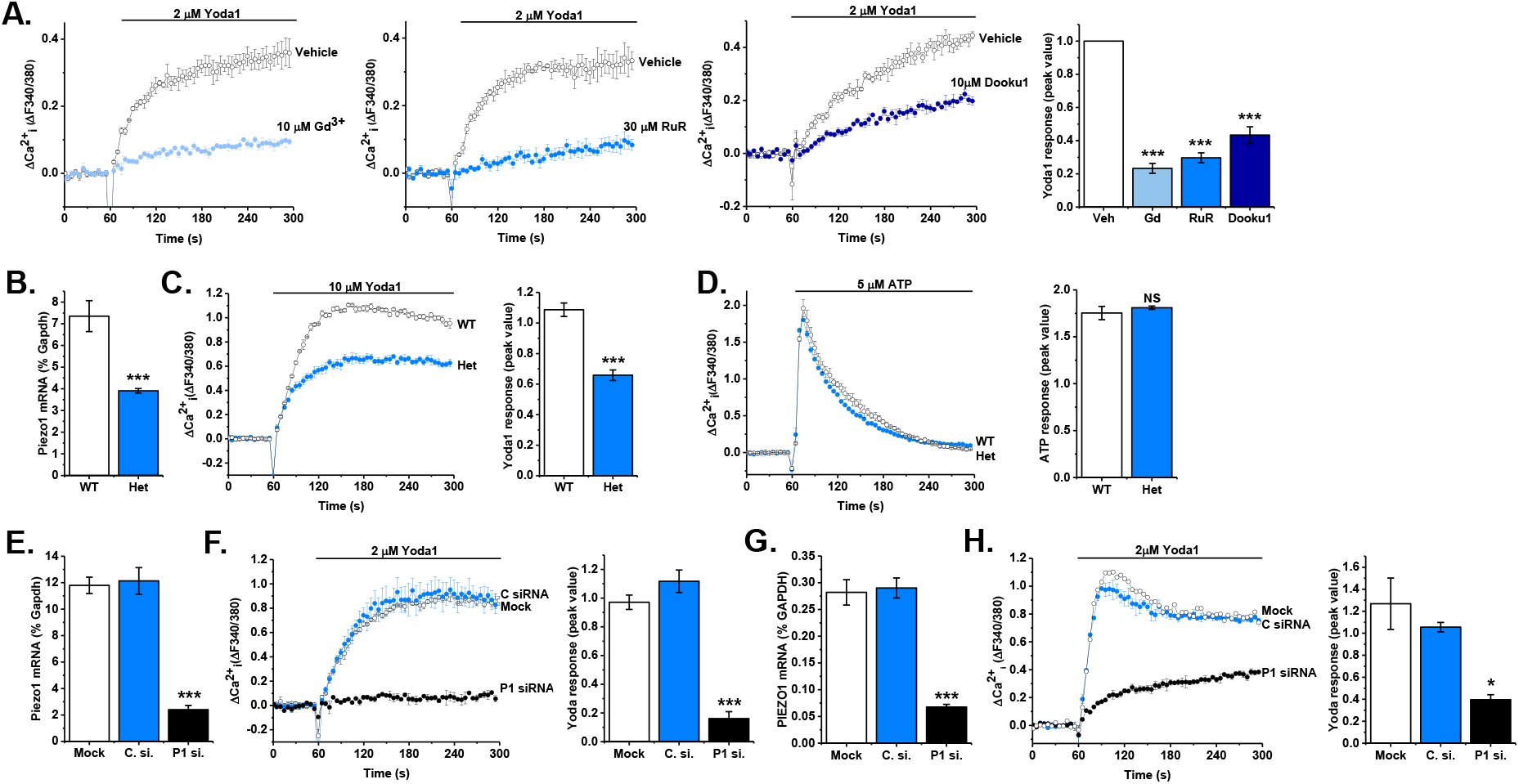
Yoda1-evoked Ca^2+^ entry is dependent on Piezo1 expression. **(A)** Representative intracellular Ca^2+^ traces and mean data after murine cardiac fibroblasts were exposed to 10 μM gadolinium (Gd^3+^), 30 μM ruthenium red (RuR), 10 μM Dooku1 or vehicle for 30 min before activation of Piezo1 by application of 2 μM Yoda1. Data was normalized to vehicle-treated cells. ***P<0.001 versus vehicle-treated cells (n/N=3/9). **(B)** RT-PCR analysis of *Piezo1* mRNA expression in cultured murine cardiac fibroblasts isolated from wild-type (WT) and *Piezo1*^+/−^ (Het) mice. Expression is measured as % of housekeeping control, *Gapdh*. ***P<0.001 (n=8). **(C)** Representative Ca^2+^ trace and mean data illustrating Ca^2+^ entry elicited by 10 μM Yoda in cardiac fibroblasts from WT (n/N=8/24) and *Piezo1*^+/−^ Het (n/N=5/15) mice. ***P<0.001. **(D)** Representative Ca^2+^ trace and mean data illustrating the Ca^2+^ entry evoked by 5 μM ATP in cardiac fibroblasts from wild-type (WT; n/N=4/12) and *Piezo1*^+/−^ (Het; n/N=3/9) mice. Not significant (NS). **(E)** RT-PCR analysis of *Piezo1* mRNA expression following transfection of murine cardiac fibroblasts with Piezo1 siRNA, mock-transfected cells and cells treated with control siRNA. Expression is measured as % of housekeeping control, *Gapdh*. ***P<0.001 versus mock-transfected cells (n=3). **(F)** Representative Ca^2+^ trace and mean data showing response to 2 μM Yoda1 in murine cardiac fibroblasts transfected with Piezo1-specific siRNA, compared to mock-transfected cells and cells treated with control siRNA. ***P<0.001 versus mock-transfected cells (n/N=3/9). **(G)** As for *E* but in human cardiac fibroblasts (n=3). **(H)** As for *F* but in human cardiac fibroblasts. *P<0.05 versus mock-transfected cells (n/N=3/9).

Together, these data established that Yoda1-induced Ca^2+^ entry in cardiac fibroblasts is dependent on Piezo1 expression and that the channel has the expected pharmacological properties.

### Activation of Piezo1 in cardiac fibroblasts opposes myofibroblast differentiation

To gain insight into the functional role of Piezo1 activation in cardiac fibroblasts, the effect of 24 h treatment with 0.5-10 μM Yoda1 on the expression of remodeling genes in murine fibroblasts was investigated by RT-PCR (**Fig. 3**). Prolonged Yoda1 treatment did not modulate *Piezo1* gene expression (**Fig. 3A**), nor did it affect expression of genes involved in ECM turnover, including collagens I and III (*Col1a1, Col3a1*) or matrix metalloproteinases (MMPs) 3 and 9 (*Mmp3, Mmp9*) (**Fig. 3B-E**). However, a concentration-dependent increase in mRNA expression of the inflammatory/hypertrophic cytokine IL-6 was observed; with significant 2 to 4-fold increases observed in response to 2-10 μM Yoda1 (**Fig. 3F**). This was not a generic inflammatory response since *Il1b* mRNA levels remained unaffected by Yoda1 treatment (**Fig. 3G**). In contrast, mRNA levels of *Acta2*, the gene that encodes α-smooth muscle actin (α-SMA), showed a decreasing trend with increasing Yoda1 concentration (**Fig. 3H**), and expression of the profibrotic cytokine transforming growth factor (TGF)-β (*Tgfb1*) was significantly reduced in a concentration-dependent manner (**Fig. 3I**).

**Figure 3.**
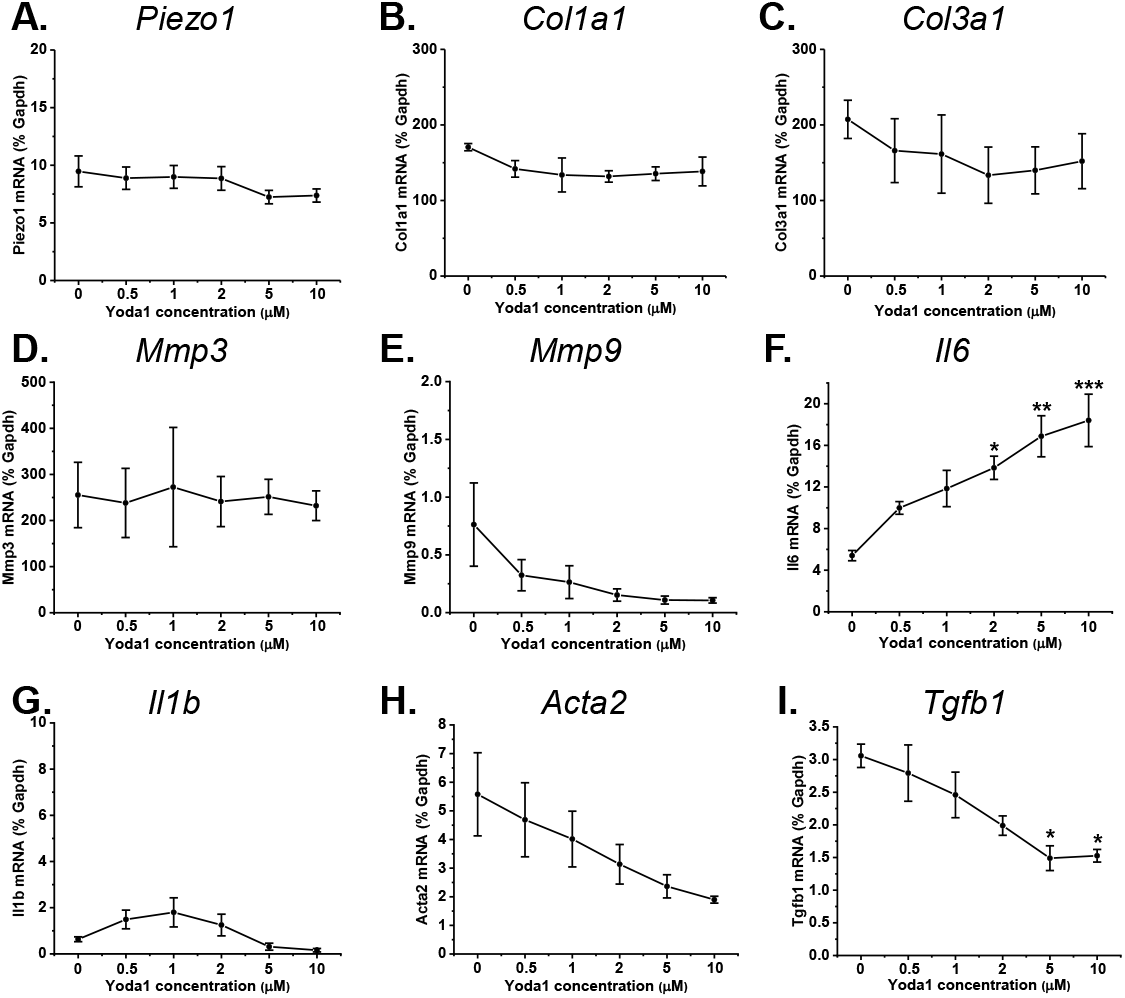
Effect of Yoda1 on gene expression in cardiac fibroblasts. **(A-I)** RT-PCR analysis of murine cardiac fibroblasts treated with concentrations of Yoda1 ranging from 0.5-10 μM, or DMSO vehicle, for 24 h. All mRNA expression levels were normalized to those of the housekeeping gene, *Gapdh*. *P<0.05, **P<0.01, ***P<0.001 versus vehicle-treated cells (n=3).

As both TGF-β and α-SMA expression are increased with fibroblast-to-myofibroblast differentiation, these gene expression data indicated that Piezo1 activation was opposing myofibroblast differentiation. Prolonged exposure to Yoda1 was also found to induce a morphological change in the cells towards a more spindle-like, less rhomboid shape, without affecting cell viability; also indicative of a less differentiated state (**Fig. 4A**). To investigate this further, collagen gel contraction assays were employed, in which increased gel contraction correlates with increased myofibroblast activity. Collagen gels seeded with mouse cardiac fibroblasts were treated with Yoda1 or with cytokines that promote the myofibroblast and fibroblast phenotypes respectively; namely TGF-β1 or IL-1α. As expected, TGF-β1 increased myofibroblast differentiation and gel contraction, resulting in a reduction in gel weight compared to both Yoda1-treated and untreated gels (**Fig. 4B**), whereas IL-1α had the opposite effect. Yoda1 stimulated a significant increase in collagen gel weight of 25% compared to unstimulated gels (**Fig. 4B**), indicating a reduction of myofibroblast phenotype in a manner similar to IL-1α. This Yoda1-evoked loss of myofibroblast phenotype was dependent upon Piezo1, as evidenced by attenuation of the Yoda1-induced response when cardiac fibroblasts were transfected with Piezo1-specific siRNA, compared to mock-transfected or control siRNA-transfected cardiac fibroblasts (**Fig. 4C**).

**Figure 4.**
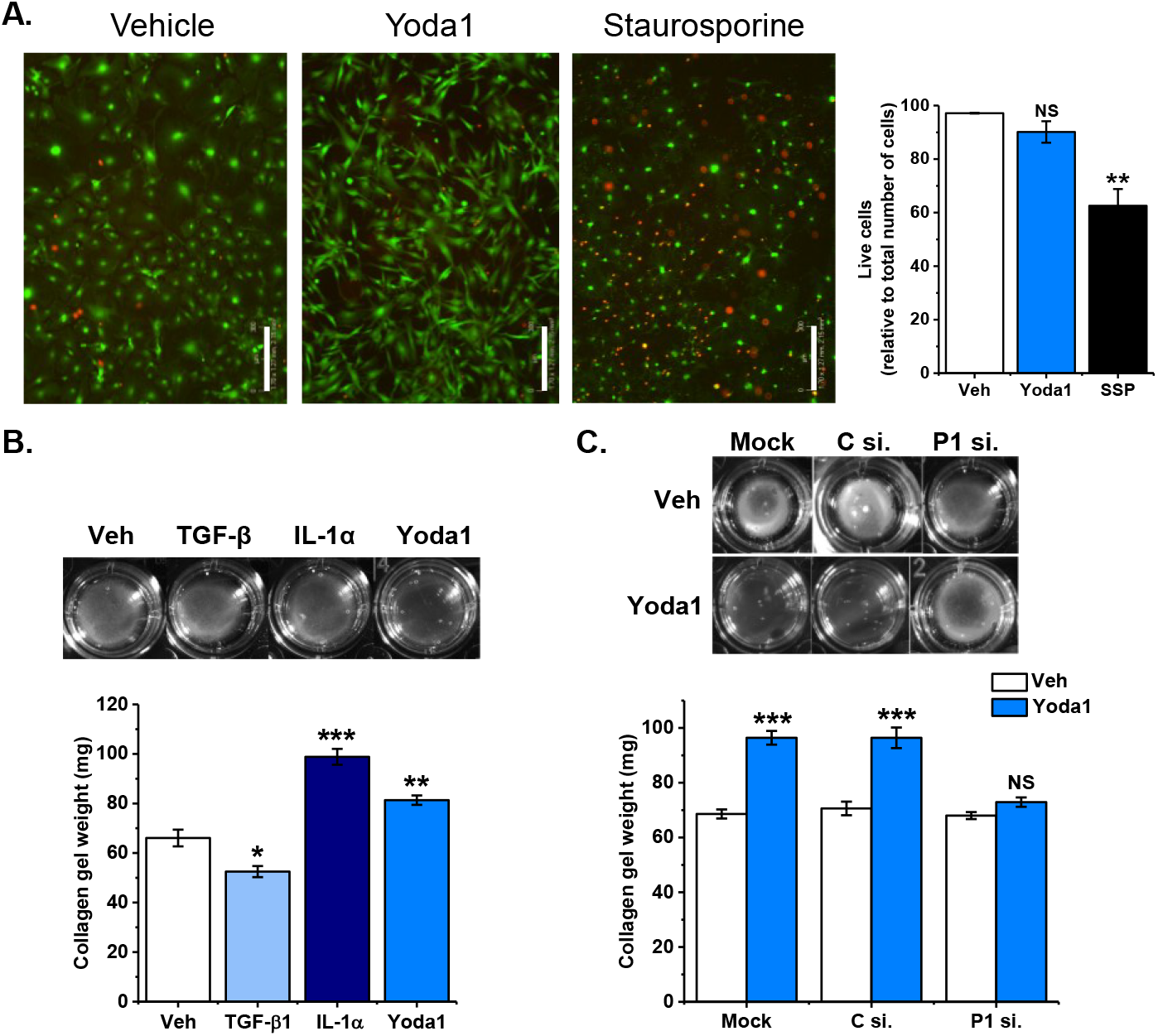
Activation of Piezo1 in cardiac fibroblasts opposes myofibroblast differentiation. **(A)** Representative images from live/dead cell assay performed on cultured murine cardiac fibroblasts treated with either vehicle, 10 μM Yoda1 or 1 μM staurosporine (SSP) for 24 h. Green indicates live cells; red indicates dead cells. Scale bar = 300 μm. Bar chart shows mean data for viable cells as a percentage of total cells. **P<0.005, not significant (NS) versus vehicle-treated cells (n=3). **(B)** Representative images of contracted gels after 24 h treatment with 1 ng/ml TGFβ, 10 ng/ml IL-1 α and 5 μM Yoda1 on collagen gels containing cultured murine cardiac fibroblasts. Collagen gel weight after 24 h treatment is shown. *P<0.05, **P<0.01, ***p<0.001 versus vehicle-treated cells (n=5). **(C)** Representative images of contracted gels after 24 hours treatment with vehicle or 5 μM Yoda1 on collagen gels containing cultured murine cardiac fibroblasts which have been either mock-transfected, transfected with control siRNA or Piezo1 siRNA. Collagen gel weight after 24 h treatment is shown. ***P<0.001, not significant (NS) versus vehicle-treated cells (n=5).

### Piezo1 activation stimulates IL-6 expression and secretion via p38 MAP kinase

Given our recent findings on the potential importance of cardiac fibroblast-derived IL-6 in modulating cardiac hypertrophy [14], and the causative link between mechanical stimulation and cardiac hypertrophy [16], we proceeded to interrogate the mechanism by which Piezo1 activation was coupled to IL-6 expression. Firstly, we explored whether mechanical activation of cardiac fibroblasts was coupled to IL-6 expression and whether this occurred via a Piezo1-dependent mechanism. Cyclic stretching (1 Hz, 10% stretch) of human cardiac fibroblasts for 6 h induced a marked increase in *IL6* mRNA expression, which was attenuated in PIEZO1 gene-silenced cells (**Fig. 5A**), indicating that activation of Piezo1 by either mechanical (stretching) or chemical (Yoda1) methods resulted in increased IL-6 expression in cardiac fibroblasts. We then probed the mechanism further in murine cardiac fibroblasts using Yoda1 as a stimulus. *Il6* mRNA expression increased in a time-dependent manner over a 2-24 h period of Yoda1 treatment (**Fig. 5B**). This increase correlated with an increase in IL-6 protein secretion (**Fig. 5C**). Yoda-1 induced IL-6 secretion was reduced by approximately 50% in cardiac fibroblasts isolated from hearts of *Piezo1*^+/−^ mice compared with those from WT hearts (**Fig. 5D**), in keeping with the reduction in Piezo1 expression and Ca^2+^ response in these cells (**Fig. 2B,C**). IL-6 secretion was unchanged following treatment of cardiac fibroblasts with compound ‘2e’ (**Fig. 5D**), an inactive analogue of Yoda1 [15] that did not induce Ca^2+^ entry in murine or human cardiac fibroblasts or in HEK T-REx-293 cells heterologously expressing mouse Piezo1 (**Suppl. Fig. 2A-C**). The Yoda1-induced increase in IL-6 secretion was dependent on Piezo1, as evidenced by attenuation of the secretion in murine cardiac fibroblasts transfected with Piezo1-specific siRNA (**Fig. 5E**). The same was true in human cardiac fibroblasts, where the Yoda1-evoked increase in *Il6* mRNA expression was attenuated by Piezo1-specific siRNA, and again the inactive Yoda1 analogue 2e had no effect (**Suppl. Fig. 2D,E**). These data establish that Piezo1 activation using either chemical or mechanical stimuli induces IL-6 expression in cardiac fibroblasts via a Piezo1-dependent mechanism.

**Figure 5.**
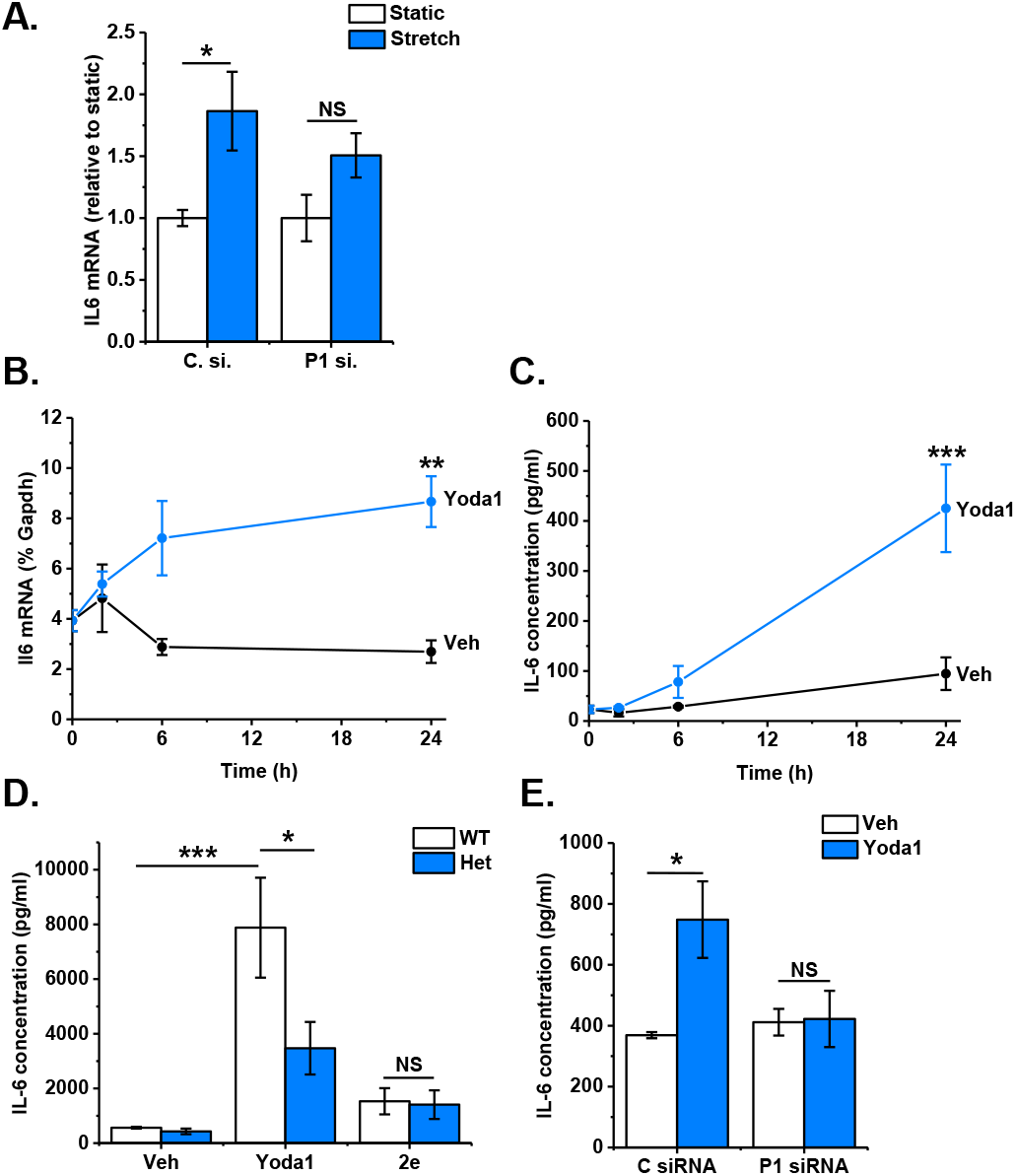
Mechanical or chemical activation of Piezo1 is coupled to IL-6 expression. **(A)** RT-PCR analysis of *IL6* mRNA expression after exposure of human cardiac fibroblasts to 6 h cyclical stretch (1 Hz, 10% stretch) for 6 h, compared to fibroblasts maintained in parallel under static conditions. Cardiac fibroblasts were previously transfected with either Piezo1-specific siRNA or control siRNA. Expression is measured as % of housekeeping control, *GAPDH* and data are normalized to static control. *P<0.05, not significant (NS) versus static cells (n=7). **(B,C)** Murine cardiac fibroblasts were exposed to vehicle or 10 μM Yoda1 for 2-24 h before (*B*) measuring *Il6* mRNA levels by RT-PCR where expression is measured as % of housekeeping control, *Gapdh* or (*C*) analyzing conditioned medium for IL-6 levels by ELISA. **P<0.01, ***P<0.001 versus vehicle-treated cells (n=3). **(D)** Cardiac fibroblasts from wild-type (WT; n=4) and *Piezo1*^+/−^ (Het; n=3) mice were treated with vehicle, 10 μM Yoda1 or compound 2e for 24 h before measuring IL-6 levels in conditioned medium using ELISA. *P<0.05, ***P<0.001, not significant (NS). **(E)** Cells were transfected with control or Piezo1-specific siRNA and treated with either vehicle or 10 μM Yoda1 for 24 h before collecting conditioned media and measuring IL-6 levels by ELISA. *P<0.05, not significant (NS) versus-vehicle-treated cells (n=3).

PamChip multiplex serine/threonine kinase activity profiling was used to assess differences in kinase activity following treatment of murine cardiac fibroblasts with Yoda1 for 10 min. Combinatorial analysis of phosphorylation of 140 peptide substrates revealed a hierarchical list of predicted serine/threonine kinases that were activated downstream of Piezo1 (**Fig. 6A, Suppl. Table 1**). Within the top 20 hits, two major kinase families were identified; MAP kinases (extracellular signal-regulated kinases ERK1/2/5, c-Jun N-terminal kinases JNK1/2/3, p38 mitogen-activated protein kinases p38α/β/γ/δ) and cyclin-dependent kinases (CDKs1-7,9,11) (**Fig. 6A**). The primary role of CDK-family kinases is to phosphorylate cell cycle proteins and thereby regulate cell cycle progression, although it has also been reported that specific CDKs can regulate inflammatory gene expression through interaction with the NF-kB pathway [17]. However, given that kinases within the NF-kB pathway (IKKα, IKKβ) were not activated in response to Yoda1 in our experiments (**Suppl. Table 1**), we did not pursue the role of CDKs further. Subsequent experiments were therefore undertaken with selective pharmacological inhibitors of the ERK (PD98059), JNK (SP600125) and p38 MAPK (SB203580) pathways to identify which ones were required for expression of IL-6 in response to Piezo1 activation in murine cardiac fibroblasts (**Fig. 6B-D**). Of these, only SB203580 significantly reduced the Yoda1-induced increase in *Il6* mRNA expression back to basal levels (**Fig. 6D**), suggesting an important role for the p38 MAPK pathway in Yoda1-induced *Il6* gene expression. In agreement, SB203580 also inhibited Yoda1-induced IL-6 protein secretion by 80% (**Fig. 6E**).

**Figure 6.**
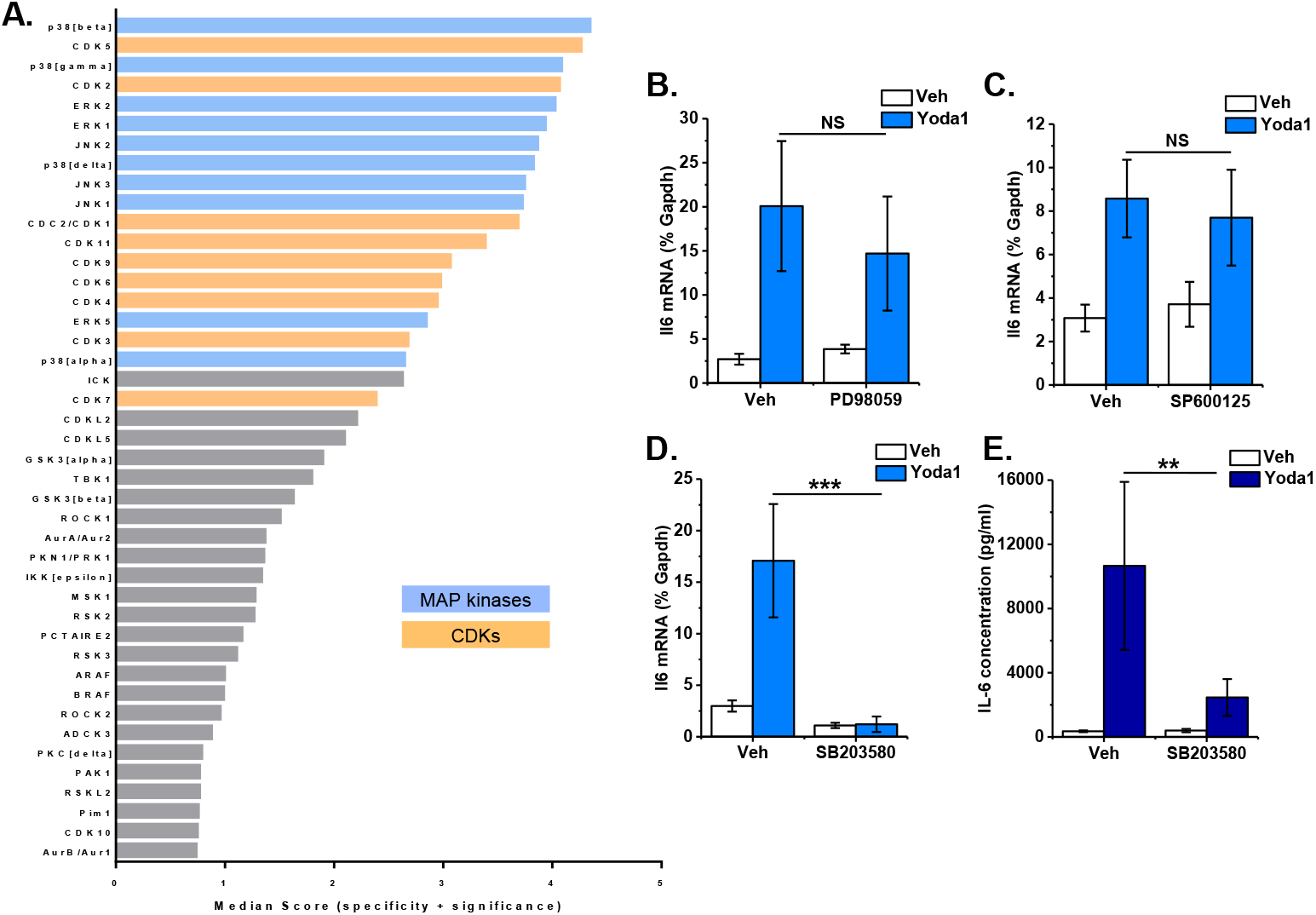
Piezo1 activation induces MAP kinase signaling with p38 MAPK being coupled to IL-6 expression. **(A)** Mouse cardiac fibroblasts were stimulated with 10 μM Yoda1 for 10 min before analysing Ser/Thr protein kinase activity with PamChip multiplex kinase activity profiling (n=6). Kinases identified based on peptide substrate phosphorylation and ranked by specificity and statistical significance (only the top 40 are shown). See Suppl. Table 1 for full data set. The top kinase families predicted to be activated by Yoda1 were the MAP kinases (ERK1/2/5, JNK1/2/3, p38α/β/γ/δ; blue) and the cyclin-dependent kinases (CDKs1-7,9,11; orange). **(B-D)** RT-PCR analysis of *Il6* mRNA expression after exposure of murine cardiac fibroblasts to (*B*) 30 μM PD98059 (ERK pathway inhibitor), (*C*) 10 μM SP600125 (JNK inhibitor) or (*D*) 10 μM SB203580 (p38 MAPK inhibitor) for 1 h followed by treatment with vehicle or 10 μM Yoda1 for a further 24 h. Expression is measured as % of housekeeping control, *Gapdh*. ***P<0.001, not significant (NS) for effect of inhibitor (n=7 for SB203580, n=5 for PD98059 and SP600125). **(E)** Murine cardiac fibroblasts were treated with vehicle or 10 μM SB203580 for 1 h and then treated with either vehicle or 10 μM Yoda1 for 24 h before collecting conditioned media and measuring IL-6 levels by ELISA. **P<0.01 for effect of inhibitor (n=6).

Western blotting was used to investigate whether the p38 MAPK pathway was stimulated following Piezo1 activation. Yoda1 increased p38 phosphorylation (activation) after 10 min before returning to basal levels (**Fig. 7A**). Further studies investigating p38 activation revealed that Yoda1-induced p38 phosphorylation occurred in a concentration-dependent manner and that p38 was not activated in response to the inactive Yoda1 analogue 2e (**Fig. 7B**). Piezo1-specific siRNA reduced Yoda1-induced activation of p38α in both murine (**Fig. 7C**) and human (**Fig. 7D**) cardiac fibroblasts, confirming the role of Piezo1 in Yoda1-induced p38 activation. The p38 inhibitor SB203580 significantly reduced the Yoda1-induced increase in p38 phosphorylation (**Fig. 7E**). Furthermore, p38 activation following Yoda1 treatment was shown to be dependent on the presence of extracellular Ca^2+^ (**Fig. 7F**), indicating that it is Ca^2+^ entry through the Piezo1 channel that is important for downstream signaling. Together, these data demonstrate that Ca^2+^-induced p38 MAPK activation is essential for inducing the expression and secretion of IL-6 in response to Piezo1 activation.

**Figure 7.**
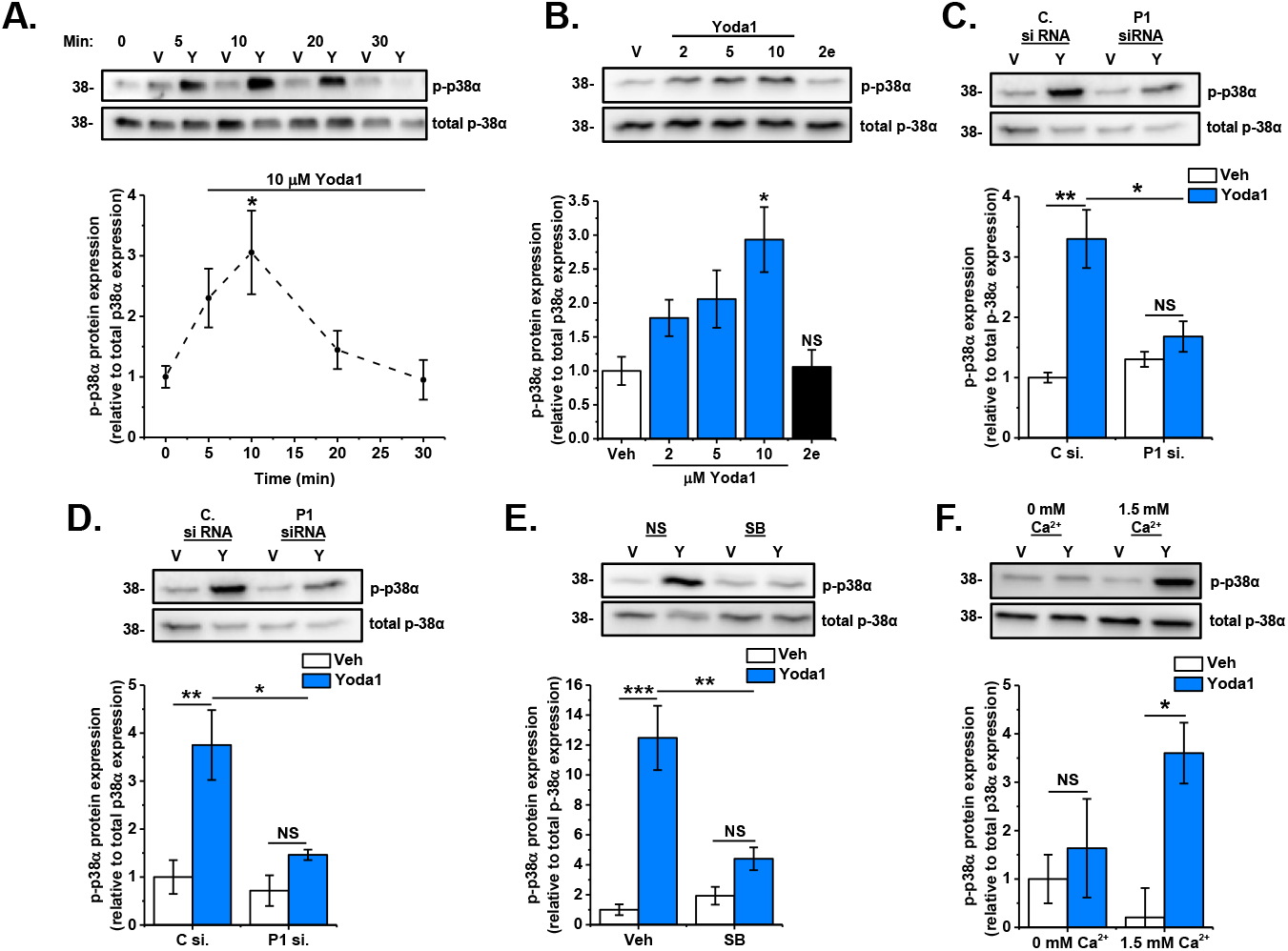
p38 MAPK phosphorylation following Yoda1 treatment is dependent on Piezo1. **(A)** Murine cardiac fibroblasts were treated with DMSO vehicle (V) or 10 μM Yoda1 (Y) for 5-30 min. Samples were immunoblotted for p-p38α expression and reprobed with total p38 antibody to confirm equal protein loading. Graph shows mean densitometric data. *P<0.05 versus vehicle-treated cells (n=6). **(B)** Murine cardiac fibroblasts were treated for 10 min with varying concentrations of Yoda1 (2-10 μM) or 10 μM compound 2e. Samples were immunoblotted for p-p38α and reprobed with antibody for total p38α, to confirm equal protein loading. Graphs show mean densitometric data. *P<0.05, not significant (NS) versus vehicle-treated cells (n=3). **(C,D)** Murine (*C*) or human (*D*) cardiac fibroblasts were transfected with either scrambled or Piezo1-specific siRNA before treatment with vehicle or 10 μM Yoda1 for 10 min. Samples were immunoblotted for p-p38α and reprobed with total p38α antibody to confirm equal protein loading. Graphs show mean densitometric data. *P<0.05, **P<0.01, not significant (NS) (n=3). **(E)** Murine cardiac fibroblasts were exposed to 10 μM SB203580 for 1 hour before treatment with vehicle or 10 μM Yoda1 for 10 min. Samples were immunoblotted for p-p38α and reprobed with total p38α, to confirm equal protein loading. Graphs show mean densitometric data. **P<0.01, ***P<0.001, not significant (NS) (n=3). **(F)** Murine cardiac fibroblasts were treated for 10 min with vehicle or 10 μM Yoda1 in either standard DMEM or in DMEM containing 1.75 mM EGTA to chelate free Ca^2+^. Samples were immunoblotted for p-p38α and reprobed with total p38α antibody to confirm equal protein loading. Graphs show mean densitometric data. *P<0.05, not significant (NS) (n=3).

## Discussion

The three main findings of this study are that (i) human and mouse cardiac fibroblasts express functional Piezo1 channels coupled to a rapid rise in intracellular Ca^2+^; (ii) Piezo1 activation opposes cardiac myofibroblast activation; and (iii) Piezo1 activation is coupled to secretion of IL-6 via a p38 MAPK-dependent pathway. These novel data establish a link between cardiac fibroblast Piezo1, myofibroblast differentiation and secretion of paracrine signaling molecules that can modulate cardiac remodeling.

We established that murine and human cardiac fibroblasts express *Piezo1* mRNA at levels similar to those found in endothelial cells from various sources, which are known to express high levels of functional Piezo1 [13]. Piezo1 mRNA expression was 20 times higher in cardiac fibroblasts than cardiomyocytes. The EC_50_ values of Yoda1 acting on endogenous Piezo1 in murine and human cardiac fibroblasts (0.72 μM and 0.71 μM respectively) were comparable with our previous report of 0.23 μM in HUVECs [15]. When artificially over-expressed in cell lines, our data on mouse Piezo1 in HEK T-REx-293 cells (EC_50_ = 0.33 μM) are broadly comparable with human Piezo1 expressed in HEK T-REx-293 cells (EC_50_ = 2.51 μM) [15], and somewhat lower than those originally reported in HEK293T cells transiently transfected with mouse (EC_50_ = 17.1 μM) or human (EC_50_ = 26.6 μM) Piezo1 [11]. It is worth noting that we could only use Yoda1 at concentrations up to 10 μM due to solubility problems, so these EC_50_ values are estimates.

The ability of common inhibitors of mechanosensitive ion channels, namely gadolinium and ruthenium red [10], and the novel Yoda1 antagonist Dooku1 [15], to decrease the Yoda1-evoked Ca^2+^ entry in murine and human cardiac fibroblasts correlated with the expected properties of the channel. The mechanism by which Yoda1 modulates Piezo1 activation and gating has been described recently [18]. However, it has also been suggested that Yoda1 may have some Piezo1-independent effects in endothelial cells [19]. We further verified the importance of Piezo1 by showing that the Ca^2+^, myofibroblast differentiation, p38 MAPK and IL-6 responses in cardiac fibroblasts were all proportionally reduced by siRNA-mediated knockdown of Piezo1 or by use of cells from a global *Piezo1*^+/−^ heterozygous knockout mouse model [13]. Thus, the Yoda1 responses that we observed in cardiac fibroblasts were unequivocally due to activation of Piezo1.

Exposure to Yoda1 for 24 h modulated expression of several genes in murine cardiac fibroblasts in a concentration-dependent manner, including increasing *Il6* expression and decreasing *Acta2* (i.e. α-SMA) and *Tgfb1*. The decrease in *Acta2* and *Tgfb1* expression is of interest as these two molecules are key markers of myofibroblast differentiation, and our data suggest that Piezo1 activation is opposing this differentiation. Further evidence for an effect on the myofibroblast phenotype came from florescence microscopy showing Yoda1 induced a morphological change in the cells towards a more spindle-like shape, similar to that which we observed in human cardiac fibroblasts after treatment with IL-1 [20]. In that study, we also showed that morphological changes induced by IL-1 were due to a reduced level of myofibroblast differentiation, as evidenced by an impaired capacity to contract collagen gels [20]. Our current data reveal that Yoda1 treatment reduced collagen gel contraction, and that this effect was dependent on Piezo1. We, and others, have shown that IL-1 is a potent inducer of MMP-9 and IL-1β expression [21–23], neither of which was affected by Yoda1 in the present study, suggesting that Yoda1 activation does not precisely mimic that of IL-1α. Our observation that Piezo1 opposes myofibroblast differentiation is intriguing as mechanical stimulation is known to be an important driver of myofibroblast differentiation which, along with TGF-β, induces changes in gene expression including upregulation of α-SMA. Activation of integrins, TRP channels and cytoskeletal complexes have been shown to translate mechanical signals into changes in cardiac fibroblast gene expression including those involved in myofibroblast differentiation [4,5]. However, our data suggest that Piezo1 acts to oppose these signals. More work is needed to precisely define the relative contributions of these various mechanosensing proteins in regulating myofibroblast differentiation.

Cardiac fibroblasts secrete IL-6, a pleiotropic pro-inflammatory cytokine that promotes cardiac fibroblast proliferation and fibrosis [24], as well as cardiac hypertrophy through its actions on cardiomyocytes [25]. Mechanical stretching of cardiac fibroblasts induces expression of several pro-fibrotic cytokines [1]. *IL6* is a mechanically regulated gene, and increased IL-6 secretion occurs upon skeletal muscle contraction where it is thought to play a beneficial role [26]. Fibroblasts from several sources have been shown to secrete IL-6 in response to mechanical stretch [27–29]. We showed that there is an increase in *IL6* gene expression produced in response to stretching of cardiac fibroblasts and established that this is mediated via Piezo1.

The p38 family of stress-activated MAPKs are known to play an important role in cardiac signaling and are activated in both acute and chronic cardiac pathologies [30]. Recently, a key role for p38α in regulating multiple aspects of cardiac fibroblast function has emerged. Fibroblast-specific knockout of *Mapk14*, the gene encoding p38α, revealed a critical role for this kinase in driving cardiac myofibroblast differentiation and fibrosis in response to ischemic injury or chronic neurohumoral stimulation [31]. In a similar model, we established a role for cardiac fibroblast p38α in stimulating cardiac hypertrophy after chronic β-adrenergic stimulation and uncovered a potential role for IL-6 acting as a paracrine inducer of cardiomyocyte hypertrophy in this context [14]. We have also shown an important role for p38α in cytokine-induced IL-6 expression and secretion in cultured human cardiac fibroblasts [32]. Thus, mechanical activation of p38α in cardiac fibroblasts, and subsequent secretion of IL-6 is likely to be important in the cardiac remodeling process. Interestingly, hydrostatic pressure can stimulate p38 MAPK signaling through Piezo1 activation in mesenchymal stem cells [33].

In conclusion, our study establishes that Piezo1 is a functional Ca^2+^-permeable ion channel in cardiac fibroblasts and is coupled to altered expression of genes important to the remodeling process. Specifically, activation of Piezo1 opposes cardiac fibroblast differentiation and induces expression and secretion of IL-6, a pro-hypertrophic cytokine with important roles after cardiac injury. Furthermore, we revealed that p38 MAPK is activated downstream of Piezo1-mediated Ca^2+^ entry to cause increased IL-6 secretion. Functional studies and *in vivo* investigation into the role of Piezo1 in cardiac fibroblasts are warranted to provide additional information on its role in regulating cardiac remodeling.

## Materials and Methods

### Reagents

Yoda1 (Tocris), staurosporine (Sigma), PD98059 (Merck), SP600125 (Cambridge Bioscience) and SB203580 (Merck) were all solubilized in dimethylsulfoxide (DMSO). ATP, gadolinium and ruthenium red were all obtained from Sigma-Aldrich and dissolved in H_2_O. Dooku1 and compound 2e [15] were synthesized at the University of Leeds and solubilized in DMSO.

### Mouse cardiac fibroblast culture

Adult C57BL/6J mice were euthanized according to guidelines of The UK Animals (Scientific Procedures) Act 1986. Cardiac fibroblast cultures were established from collagenase-digested hearts and cultured in Dulbecco’s Modified Eagle’s medium (DMEM) supplemented with 10% fetal calf serum, as previously described [14]. All cells were kept at 37 °C and 5% CO_2_ throughout the study. Cells were studied at passage 1-2. Cells were kept in serum-free DMEM for 16 h prior to treatment in order to collect RNA, lysates and conditioned media.

### Human cell culture

Biopsies of human atrial appendage and long saphenous vein were obtained from patients undergoing elective coronary artery bypass grafting at the Leeds General Infirmary following local ethical committee approval and informed patient consent. Cells were harvested and cultured as described previously for cardiac fibroblasts [34] and saphenous vein endothelial cells [35]. Experiments were performed on human cardiac fibroblasts from passages 2-5. Serum-free DMEM was used for 1 h prior to treatment in order to collect RNA and conditioned media. HUVECs were purchased from Lonza and cultured in Endothelial Cell Growth Medium (EGM-2) (Lonza), supplemented with 2% fetal calf serum and EGM-2 SingleQuots Kit (Lonza). HUVECs were used at passage 2.

### HEK T-Rex-293 cell culture

pcDNA3_mouse Piezo1_IRES_GFP, a kind gift from Artem Patapoutian [10], was used as a template to clone the mouse Piezo1 coding sequence into pcDNA4/TO. Overlapping mouse Piezo1 (forward primer 5’ GTAACAACTCCGCCCCATTG 3’ and reverse primer 5’ GCTTCTACTCCCTCTCACGTGTC 3’) and pcDNA4/TO (forward primer 5’ GACACGTGAGAGGGAGTAGAAGCCGCTGATCAGCCTCGACTG 3’ and reverse primer 5’ CAATGGGGCGGAGTTGTTAC 3’) PCR products were assembled using Gibson Assembly (New England Biolabs) [36]. This construct does not contain tetracycline operator sequences. HEK T-REx-293 cells (Invitrogen) were transfected with pcDNA4/TO-mPiezo1 using Lipofectamine 2000 (Invitrogen) and treated with 200 μg/ml zeocin (InvivoGen) to select for stably transfected cells. Individual clones were isolated and analyzed for expression using Yoda1 and intracellular Ca^2+^ measurements. HEK T-REx-293 cells were maintained in DMEM supplemented with 10% fetal calf serum and 1% penicillin/streptomycin (Sigma-Aldrich). Non-transfected HEK T-Rex-293 cells were used as control cells.

### Cardiac cell fractionation

Non-myocyte cardiac cell fractions were prepared as described previously [14]. Briefly, collagenase-digested heart tissue was filtered through a 30 μm MACS smart strainer (Miltenyi) to remove cardiomyocytes. Non-myocytes were then separated into two fractions using a cardiac fibroblast magnetic antibody cell separation kit (MACS; Miltenyi). ‘Non-fibroblasts’ (endothelial cells and leukocytes - *Pecam1*-positive) were collected in fraction 1, and ‘fibroblasts’ (*Col1a1/Col1a2/Ddr2/Pdgfra*-positive) were collected in fraction 2, as previously characterized [14]. Separately, adult mouse cardiomyocytes were isolated from ventricles of 8 week old WT mice. Hearts were cannulated through the aorta and perfused with perfusion buffer (124.5 mM NaCl, 10 mM HEPES, 11.1 mM glucose, 1.2 mM NaH_2_PO_4_, 1.2 mM MgSO_4_, 4mM KCl, 25 mM Taurine; pH 7.34) containing 10 mM butanedione monoxime for 5 min followed by perfusion buffer containing 1 mg/ml type 2 collagenase, 0.05 mg/ml protease and 12.5 μM CaCl_2_ for 7-15 min until suitable digestion was observed. Ventricles were gently cut in perfusion solution containing 10% FBS and 12.5 μM CaCl_2_ and filtered. This step was repeated in perfusion solution containing 5% FBS and 12.5 μM CaCl_2_ and finally perfusion solution containing only 12.5 μM CaCl_2_. The filtrate was pelleted by gravity for 5 min before resuspension in perfusion buffer containing 0.5 mM CaCl_2_. This step was repeated with increasing concentrations of CaCl_2_ (1.5 mM, 3.5 mM, 8 mM and 18 mM). Finally, the cell pellet was resuspended in 1 ml Trizol. RNA was extracted from cardiac cell fractions and qRT-PCR used to quantify Piezo1 mRNA.

### Quantitative RT-PCR

For gene expression studies, cardiac fibroblasts were treated with Yoda1 or vehicle (DMSO) for the indicated time. RNA was extracted from cultured/fractionated cells with the Aurum RNA Extraction Kit (Bio-Rad). cDNA was synthesized using a reverse-transcription system (Promega). Real-time RT-PCR was performed with the ABI-7500 System, gene expression master mix and specific Taqman primer/probe sets (Thermo Fisher Scientific): mouse *Acta2* (Mm00725412_s1), mouse *Col1a1* (Mm01302043_g1), mouse *Col3a1* (Mm01254476_m1), mouse *Piezo1* (Mm01241545_g1), human *PIEZO1* (Hs00207230_m1), mouse *Il1b* (Mm00434228_m1), mouse *Il6* (Mm00446190_m1), human *IL6* (Hs00174131_m1), mouse *Mmp3* (Mm00440295_m1), mouse *Mmp9* (Mm00442991_m1) and mouse *Tgfb1* (Mm01178820_m1). Data are expressed as percentage of mouse *Gapdh* (Mm99999915_g1) or human *GAPDH* (Hs99999905_m1) housekeeping gene mRNA expression using the formula 2^−ΔCT^ x 100, in which C_T_ is the cycle threshold number.

### Piezo1-modified mice

Animal use was authorized by both the University of Leeds Animal Ethics Committee and by The UK Home Office under project license P144DD0D6. Animals were maintained in Optimice individually ventilated cages (Animal Care Systems) at 21°C, 50-70% humidity, light/dark cycle 12 h/12 h on RM1 diet (Special Diet Services) *ad libitum* and bedding of Pure’o Cell (Datesand). C57BL/6 mice carrying global disruption of the *Piezo1* gene with a *lacZ* insertion flanked by FRT sites have been described previously [13]. Identification of wild-type (WT) and *Piezo1*^+/−^ mice was performed by genotyping using primers for *LacZ*: forward: AATGGTCTGCTGCTGCTGAAC and reverse: GGCTTCATCCACCACATACAG. Mice of varying ages and sexes were used for experiments.

### Western blotting

Cells were treated with vehicle (DMSO), Yoda1 or compound 2e at the appropriate times prior to lysing. Cells were harvested in lysis buffer containing 10 mM Tris, pH 7.5, 150 mM NaCl, 0.5 mM EDTA, 0.5 % NP-40, MiniComplete protease inhibitors (Roche), and PhosSTOP phosphatase inhibitors (Roche). A protein quantification assay was then performed using the DC Protein Assay (BioRad). 25 μg protein was loaded on a 10% polyacrylamide gel. After resolution by electrophoresis, samples were transferred to PVDF membranes and western blotting performed as described previously [37] using a primary antibody for phospho-p38 MAPK (#9215; Cell Signaling Technology). Membranes were re-probed with p38α antibody (#9228; Cell Signaling Technology) to confirm equal protein loading. Species-appropriate secondary antibodies (GE Healthcare) and ECL Prime Western Blotting Detection reagent (GE Healthcare) were used for visualization. Syngene G:BOX Chemi XT4 was used for imaging, alongside GeneSys image acquisition software for densitometric analysis.

### Intracellular Ca^2+^ measurements

Changes in intracellular Ca^2+^ (Ca^2+^i) concentration were measured using the ratiometric Ca^2+^ indicator dye, Fura-2-AM and a 96-well fluorescence plate reader (FlexStationII^384^, Molecular Devices), controlled by SOFTmax PRO software v5.4.5. Cardiac fibroblasts and HUVECs were plated in clear 96-well plates (Corning) and HEK T-REx-293 cells in black, clear-bottomed 96-well plates (Grenier) at a confluence of 90%, 24 h before experimentation. Cells were incubated for 1 h at 37 °C in standard bath solution (SBS: 130 mM NaCl, 5 mM KCl, 8 mM D-glucose, 10 mM HEPES, 1.2 mM MgCl_2_, 1.5 mM CaCl_2_, pH 7.4) containing 2 μM Fura-2, in the presence of 0.01% pluronic acid (Thermo Fisher Scientific) to aid dispersion. During all Fura-2 assays involving cardiac fibroblasts, 2.5 mM probenecid (Sigma-Aldrich) was present to prevent extrusion of the Fura-2 indicator [38]. Cells were washed with SBS and then incubated at room temperature for 30 min, during which inhibitors were added if applicable. Stimuli were injected at 60 sec of recording. The change in intracellular Ca^2+^ concentration (ΔCa^2+^i) was measured as the ratio of Fura-2 emission (510 nm) intensities at 340 nm and 380 nm.

### Gene silencing

Murine cardiac fibroblasts were grown to 80% confluence and transfected with 10 nM Piezo1-specific Silencer Select Pre-Designed siRNA (4390771, siRNA ID: s107968, Life Technologies) or Silencer Select Negative Control No. 1 siRNA (4390843, Life Technologies), using Lipofectamine RNAiMAX Reagent (Life Technologies) in OptiMEM (Gibco), as per the manufacturer’s instructions. Medium was replaced with full-growth media 24 h later. For human cardiac fibroblasts, cells were grown to 90% confluence and transfected with 20 nM Piezo1-specific Silencer Select Pre-designed siRNA (#4392420, siRNA ID: s18891, Thermo Fisher Scientific) or ON-TARGETplus Non-targeting Pool siRNA (#D-01810-10-20, Dharmacon) using Lipofectamine 2000 in OptiMEM, as per the manufacturer’s instructions. Medium was replaced with full-growth medium after 4.5 h. Cells were used for experimentation 48 h after transfection.

### ELISA

Cells were pre-treated for 24 h with vehicle (DMSO), Yoda1 or compound 2e. Conditioned media were collected, centrifuged to remove cellular debris and stored at −20 °C for subsequent analysis. The concentration of IL-6 in media was measured by ELISA (M6000, R & D Systems), according to the manufacturer’s instructions. Samples were diluted 1:5-1:30 prior to analysis.

### Stretch experiments

Human cardiac fibroblasts were seeded at 1×10^5^ cells/well on collagen-coated membranes (BioFlex 6-well culture plates). 72 h after transfection, cells were stretched whilst in serum-free medium using an FX-4000 or FX-5000 Flexercell Tension System (Flexcell International) to equibiaxially elongate the cell-seeded elastic membrane against a loading post. Elongation at 10% strain and 1 Hz was applied to the cells for 6 h. A 6-well stretching plate was housed inside the incubator (37°C, 5% CO_2_) alongside unstimulated cells adhered to Bioflex plates which served as static controls. RNA was extracted and qRT-PCR used to quantify gene expression.

### Collagen gel contraction assay

The collagen gel contraction assay was performed as described previously in 24-well plates [20]. Wells were coated with bovine serum albumin (Sigma) at 37 °C for 1 h. Collagen gels containing cells were prepared by mixing type I rat tail collagen solution (Merck) with 2 x concentrated DMEM (Merck), supplemented with 88 mM NaHCO_3_, and then immediately mixing with freshly trypsinized cells. Gels were allowed to solidify at 37 °C for 1 h. Following solidification, gels were released from the sides of the well using a pipette tip and 0.5 ml DMEM was added to the gels, along with Yoda1, TGF-β1 or IL-1α (5 μM, 1 ng/ml and 10 ng/ml respectively). Gels were photographed and weighed 24 h later to assess their relative contraction.

### Cell viability assay

Cardiac fibroblasts were plated at 80% confluency and incubated overnight (37°C, 5% CO_2_). Vehicle (DMSO), Yoda1 (10 μM) or staurosporine (1 μM) were applied the following day and cells were incubated for 24 h prior to commencing the LIVE/DEAD cell viability assay (ThermoFisher Scientific), according to the manufacturer’s instructions. Cells were imaged using the IncuCyte ZOOM Live Imaging System (Essen Bioscience). The total number of fluorescent cells in each well was calculated using inbuilt algorithms, using an average from 9 images per well of a 12 well plate. The mean data are shown as the number of live cells relative to the total number of cells.

### Multiplex kinase activity profiling

PamGene Serine-Threonine Kinase (STK) multiplex activity assays were used to investigate STK activity. Murine cardiac fibroblasts were treated with vehicle or Yoda1 for 10 min, collected and lyzed using the M-PER lysis buffer (Thermo Fisher Scientific) containing Halt phosphatase and Halt protease inhibitor cocktails (Pierce) for 30 min on ice. A protein quantification assay was then performed as stated above and samples were snap-frozen in liquid nitrogen. 1.2 μg protein was loaded on to each STK PamChip array. The phosphorylation of PamChip peptides were measured by the PamStation 12 (Hertogenbosch, Netherlands), following the manufacturer’s protocols described previously [39]. Signal intensities were analyzed using PamGene’s BioNavigator software comparing Yoda1 treatment with DMSO treatment after 10 min. Permutation analysis resulted in a specificity score (mapping of peptides to kinases) and a significance score (difference between treatment groups) for each kinase. The combination of these two scores was used to rank and predict top kinase hits.

### Statistical analysis

OriginPro 2015 (OriginLab) was used for data analysis and presentation. Averaged data are presented as mean ± SEM, where n represents the number of independent experiments and N indicates the total number of replicates within the independent experiments. For comparisons between two sets of data, a Student’s t-test was used. For multiple comparisons, a one-way ANOVA was used with Tukey post-hoc test or Sidak post-hoc test if there were differing numbers of independent experiments. P<0.05 was considered statistically significant. For IC_50_ determination, data were normalized to vehicle and curves were fitted using the Hill1 equation.

## Abbreviations

α-SMA,: α-smooth muscle actin;
CDK,: cyclin-dependent kinase;
DMSO,: dimethylsulfoxide;
ECM,: extracellular matrix;
ERK,: extracellular signal-regulated kinases;
Gapdh/GAPDH,: glyceraldehyde-3-phosphate dehydrogenase;
HEK,: human embryonic kidney;
Het,: heterozygous;
HUVECs,: human umbilical vein endothelial cells;
IL,: interleukin;
JNK,: c-Jun N-terminal kinase;
MACS,: magnetic antibody cell separation;
MAPK,: mitogen-activated protein kinase;
MMP,: matrix metalloproteinase;
SBS,: standard bath solution;
STK,: Serine-Threonine Kinase;
WT,: wild-type.

## Acknowledgments

We are grateful to Artem Patapoutian (Scripps Research Institute, CA, USA) for provision of the pcDNA3_mouse Piezo1_IRES_GFP plasmid. We gratefully acknowledge Savithri Rangarajan (PamGene) for assistance in analysis of data from PamChip kinase activity arrays. We are grateful to David Eisner’s group at the University of Manchester for supplying the mouse cardiomyocyte fraction.

## Author contributions

NMB, VS, MJL, HTJG, ELE and KC performed the research. NAT, DJB and JL obtained funding for the research. NAT, DJB, MJD, KEP, FAvN, JS and RF supervised the research. NMB and VS analyzed the data. NMB and NAT wrote the manuscript.

## Data and materials availability

All data needed to evaluate the conclusions in the paper are present in the paper or the Supplementary Materials.

**Supplementary Figure 1.**
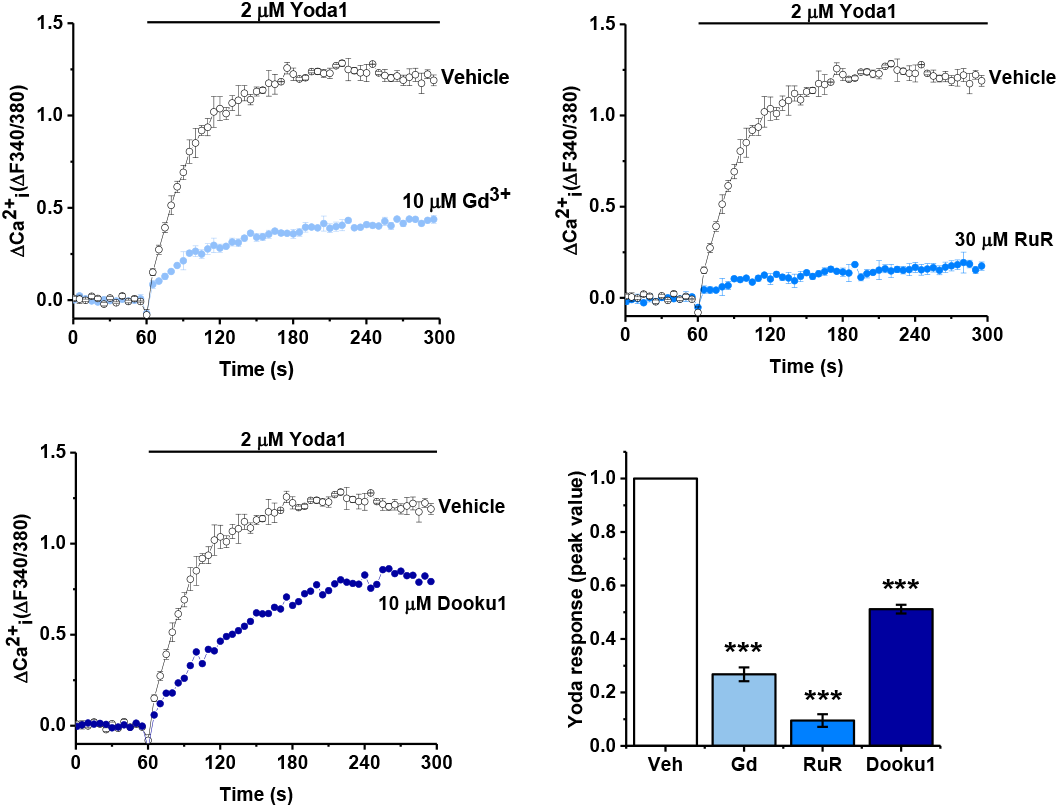
Inhibition of Yoda1-evoked Ca^2+^ entry in human cardiac fibroblasts. Representative intracellular Ca^2+^ traces and mean data after human cardiac fibroblasts were exposed to 10 μM gadolinium (Gd^3+^), 30 μM ruthenium red (RuR), 10 μM Dooku1 or vehicle for 30 min before activation of Piezo1 by application of 2 μM Yoda1. Data was normalized to vehicle-treated cells. ***P<0.001 versus vehicle-treated cells (n/N=3/9).

**Supplementary Figure 2.**
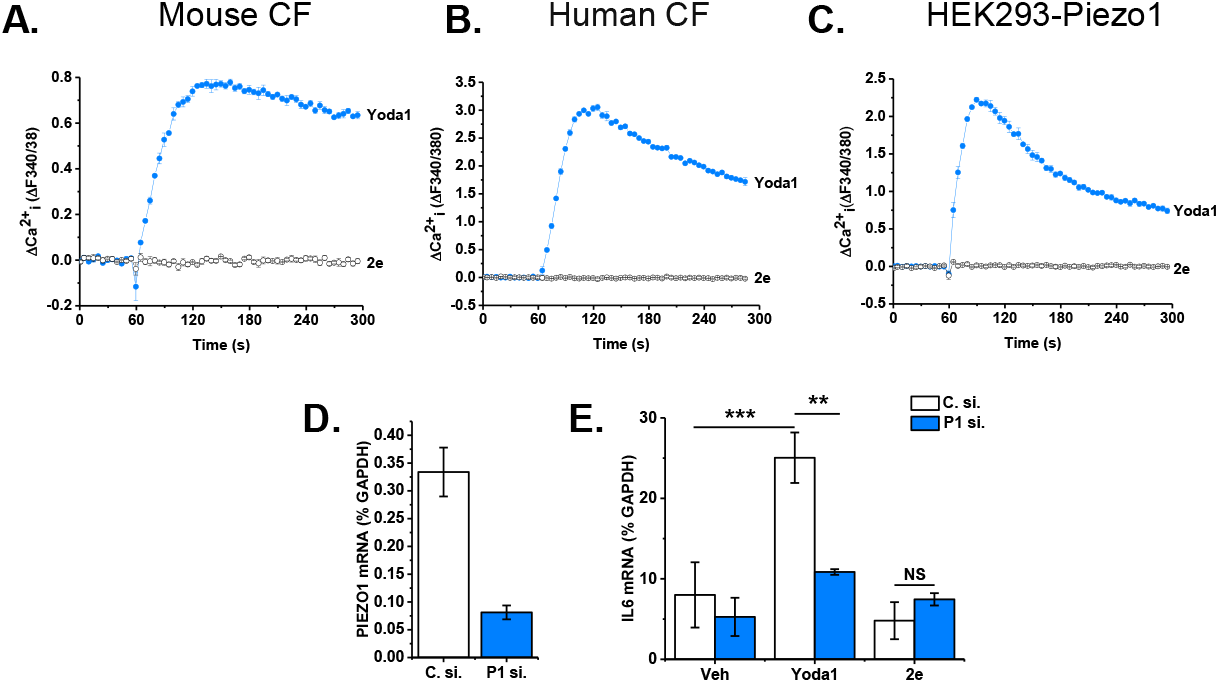
Yoda1, but not compound 2e, induces Ca^2+^ entry and IL-6 expression in human cardiac fibroblasts. Representative Ca^2+^ traces illustrating Ca^2+^ entry evoked by 10 μM Yoda1 or 10 μM compound 2e in **(A)** murine cardiac fibroblasts, **(B)** human cardiac fibroblasts or **(C)** HEK T-REx-293 cells heterologously expressing mouse Piezo1 (n/N=3/9). Human cardiac fibroblasts transfected with scrambled or Piezo1-specific siRNA were exposed to vehicle, 10 μM Yoda1 or compound 2e for 6 h before measuring mRNA levels by RT-PCR with primers for **(D)** *PIEZO1* or **(E)** *IL6*. Expression is measured as % of housekeeping control, *GAPDH*. **P<0.01, ***P<0.01, not significant (NS) (n=3).

**Supplementary Table 1.** PamChip multiplex kinase activity profiling of Yoda1-induced serine/threonine kinase activity. Mouse cardiac fibroblasts (n=6) were treated with 10 μM Yoda1 for 10 min before extracting cell lysates and assessing activity. Table shows kinases ranked by median score (combination of specificity and significance).

| Ranking | Kinase Name | Median Final score | Mean Specificity Score | Mean Significance Score |
| --- | --- | --- | --- | --- |
| 1 | p38[beta] | 4.36 | 2.32 | 1.75 |
| 2 | CDK5 | 4.28 | 2.47 | 1.63 |
| 3 | p38[gamma] | 4.10 | 1.97 | 1.28 |
| 4 | CDK2 | 4.08 | 2.65 | 1.39 |
| 5 | ERK2 | 4.04 | 2.70 | 1.39 |
| 6 | ERK1 | 3.95 | 2.70 | 1.30 |
| 7 | JNK2 | 3.88 | 2.70 | 1.19 |
| 8 | p38[delta] | 3.84 | 2.63 | 1.30 |
| 9 | JNK3 | 3.76 | 2.70 | 1.04 |
| 10 | JNK1 | 3.74 | 2.70 | 1.03 |
| 11 | CDC2/CDK1 | 3.70 | 2.34 | 1.24 |
| 12 | CDK11 | 3.40 | 1.78 | 1.65 |
| 13 | CDK9 | 3.08 | 1.83 | 1.27 |
| 14 | CDK6 | 2.99 | 1.55 | 1.50 |
| 15 | CDK4 | 2.96 | 1.50 | 1.44 |
| 16 | ERK5 | 2.86 | 1.74 | 1.04 |
| 17 | CDK3 | 2.69 | 1.77 | 1.18 |
| 18 | p38[alpha] | 2.66 | 1.64 | 0.93 |
| 19 | ICK | 2.64 | 1.37 | 1.26 |
| 20 | CDK7 | 2.40 | 1.44 | 0.93 |
| 21 | CDKL2 | 2.22 | 1.22 | 1.00 |
| 22 | CDKL5 | 2.11 | 1.16 | 0.94 |
| 23 | GSK3[alpha] | 1.91 | 0.93 | 0.75 |
| 24 | TBK1 | 1.81 | 1.07 | 0.94 |
| 25 | GSK3[beta] | 1.64 | 0.87 | 0.65 |
| 26 | ROCK1 | 1.52 | 0.80 | 0.71 |
| 27 | AurA/Aur2 | 1.38 | 0.72 | 0.62 |
| 28 | PKN1/PRK1 | 1.37 | 0.66 | 0.63 |
| 29 | IKK[epsilon] | 1.35 | 0.79 | 0.59 |
| 30 | MSK1 | 1.29 | 0.73 | 0.55 |
| 31 | RSK2 | 1.28 | 0.74 | 0.53 |
| 32 | PCTAIRE2 | 1.17 | 0.63 | 0.54 |
| 33 | RSK3 | 1.12 | 0.64 | 0.46 |
| 34 | ARAF | 1.01 | 0.61 | 0.41 |
| 35 | BRAF | 1.00 | 0.59 | 0.40 |
| 36 | ROCK2 | 0.97 | 0.35 | 0.43 |
| 37 | ADCK3 | 0.89 | 0.40 | 0.42 |
| 38 | PKC[delta] | 0.80 | 0.27 | 0.34 |
| 39 | RSKL2 | 0.78 | 0.43 | 0.35 |
| 40 | PAK1 | 0.78 | 0.43 | 0.35 |
| 41 | Pim1 | 0.77 | 0.51 | 0.41 |
| 42 | CDK10 | 0.76 | 0.40 | 0.39 |
| 43 | AurB/Aur1 | 0.75 | 0.42 | 0.36 |
| 44 | RSK1/p90RSK | 0.72 | 0.39 | 0.33 |
| 45 | Pim3 | 0.71 | 0.42 | 0.40 |
| 46 | RAF1 | 0.70 | 0.36 | 0.33 |
| 47 | PFTAIRE2 | 0.69 | 0.42 | 0.38 |
| 48 | PKG1 | 0.69 | 0.30 | 0.36 |
| 49 | Pim2 | 0.64 | 0.36 | 0.39 |
| 50 | DAPK3 | 0.63 | 0.30 | 0.33 |
| 51 | PKC[iota] | 0.63 | 0.23 | 0.34 |
| 52 | CDKL1 | 0.62 | 0.30 | 0.37 |
| 53 | PKC[zeta] | 0.62 | 0.29 | 0.37 |
| 54 | PKD1 | 0.60 | 0.32 | 0.28 |
| 55 | MSK2 | 0.60 | 0.30 | 0.30 |
| 56 | DCAMKL1 | 0.59 | 0.29 | 0.29 |
| 57 | PKC[beta] | 0.57 | 0.24 | 0.33 |
| 58 | CK2[alpha]1 | 0.56 | 0.33 | 0.23 |
| 59 | SGK1 | 0.56 | 0.28 | 0.27 |
| 60 | HGK/ZC1 | 0.53 | 0.23 | 0.24 |
| 61 | CHK1 | 0.51 | 0.23 | 0.30 |
| 62 | Akt2/PKB[beta] | 0.51 | 0.21 | 0.33 |
| 63 | PRKY | 0.46 | 0.29 | 0.31 |
| 64 | p70S6K | 0.45 | 0.19 | 0.32 |
| 65 | PKG2 | 0.45 | 0.15 | 0.31 |
| 66 | mTOR/FRAP | 0.45 | 0.16 | 0.21 |
| 67 | NuaK1 | 0.44 | 0.20 | 0.23 |
| 68 | IKK[beta] | 0.42 | 0.22 | 0.24 |
| 69 | Akt1/PKB[alpha] | 0.41 | 0.14 | 0.29 |
| 70 | PKC[alpha] | 0.39 | 0.10 | 0.30 |
| 71 | PKC[epsilon] | 0.35 | 0.11 | 0.28 |
| 72 | PKC[eta] | 0.32 | 0.12 | 0.28 |
| 73 | MAPKAPK2 | 0.32 | 0.05 | 0.26 |
| 74 | PKC[theta] | 0.31 | 0.05 | 0.26 |
| 75 | MAPKAPK3 | 0.30 | 0.03 | 0.26 |
| 76 | PKC[gamma] | 0.29 | 0.08 | 0.23 |
| 77 | ANP[alpha] | 0.25 | 0.04 | 0.22 |
| 78 | CK1[epsilon] | 0.24 | 0.10 | 0.14 |
| 79 | PRKX | 0.24 | 0.02 | 0.21 |
| 80 | ATR | 0.23 | 0.08 | 0.15 |
| 81 | CK1[alpha] | 0.22 | 0.12 | 0.20 |
| 82 | PKA[alpha] | 0.20 | 0.00 | 0.22 |
| 83 | p70S6K[beta] | 0.20 | 0.02 | 0.19 |
| 84 | PFTAIRE1 | 0.20 | 0.08 | 0.12 |
| 85 | CHK2 | 0.20 | 0.02 | 0.18 |
| 86 | DAPK2 | 0.19 | 0.05 | 0.13 |
| 87 | IKK[alpha] | 0.18 | 0.09 | 0.13 |
| 88 | AMPK[alpha]1 | 0.14 | 0.01 | 0.13 |
| 89 | CaMK4 | 0.13 | 0.01 | 0.12 |
| 90 | SGK2 | 0.10 | 0.00 | 0.10 |
| 91 | CaMK2[alpha] | 0.02 | 0.01 | 0.01 |

